# Early intermittent low-dose sclerostin antibody treatment promotes surface bone formation and reduces bone loss, but also decreases osteocyte apoptosis and mechanotransduction in ovariectomized rats: a pilot study

**DOI:** 10.1101/2024.10.02.616263

**Authors:** Syeda Masooma Naqvi, Wahaaj Ali, Hollie Allison, Laura M. O’Sullivan, Gill Holdsworth, Juan Alberto Panadero-Perez, Jessica Schiavi-Tritz, Laoise M. McNamara

## Abstract

Neutralizing sclerostin antibodies (Scl-Ab) mitigate bone loss and promote bone formation to address fracture risk in postmenopausal osteoporosis. Clinically, this treatment is administered monthly for women at high risk of fragility fractures, who are often years into menopause. Preclinical studies have demonstrated that dampening of bone formation occurs with continuous dosing at supraphysiological doses. Osteoporotic bone loss occurs rapidly during early menopause, followed by longer-term changes in bone mineralization and osteocyte activity. Whether earlier administration of lower-exposure Scl-Ab can mitigate bone loss and osteocyte-driven secondary mineralisation is unknown. The objective of this study was to evaluate the effects of early intermittent low-dose Scl-Abon: (1) osteoclastogenesis and bone resorption, (2) perilacunar remodelling, (3) secondary mineralization, and (4) osteocyte mechanosensitivity. Female retired breeder Wistar rats underwent bilateral ovariectomy and received monthly low-dose Scl-Ab injections (2 mg/kg/month) from 3 to 14 weeks post-OVX, while a control group remained untreated. Early intermittent low-dose Scl-Ab treatment increased bone formation and reduced osteoclastogenesis and catabolic gene expression ((Sost, Ctsk, Mmp9) compared to untreated rats. Treatment also decreased the percentage of empty lacunae and the number of MMP14+ osteocytes, accompanied by lower perilacunar mineral density and smaller lacunar size, indicating improved osteocyte survival and reduced perilacunar remodelling. Conversely, expression of osteocyte-mediated mineralization genes (DMP1, PHEX, OPN, ALP) and mechanotransduction-related genes (Vcl, integrins α5, αV, β1, CX43, Axin2, IFT88, Adcy6, Pkd1, Cav1) were reduced. Together, these findings suggest that early intermittent low-dose Scl-Ab therapy promotes surface bone formation while attenuating osteocyte-mediated secondary mineralization after initial bone loss.

**Mini Abstract:** Early intermittent low-dose sclerostin antibody treatment reduced osteoclastogenesis, bone resorption, and perilacunar remodelling, while promoting bone formation, decreasing osteocyte apoptosis, and downregulating genes associated with secondary mineralization and mechanosensitivity in ovariectomized rats. These findings suggest early intervention with Scl-Ab enhances bone formation and limits osteocyte apoptosis and subsequent secondary mineralization.

## Introduction

Postmenopausal osteoporosis is characterized by excessive bone resorption, followed by secondary bone tissue mineralization and altered osteocyte activity [1-3]. Rapid trabecular bone loss occurs during the first two years following menopause, slowing thereafter to approximately 1% per year [4]. This early phase is associated with accelerated bone turnover and structural deterioration [5, 6]. In rodent models, estrogen deficiency induces early trabecular bone loss within four weeks post-ovariectomy, followed by prolonged secondary mineralization characterized by increased matrix mineral density, microporosity, and perilacunar remodelling [1-3]. Osteocyte apoptosis and perilacunar resorption contribute to localized hypermineralization (micropetrosis) and reduced bone toughness [2, 3, 7], yet strategies to inhibit pathological mineralisation by osteocytes have not been explored.

Osteocytes synthesize and release sclerostin, a glycoprotein that plays a central role in regulating osteoclast resorption and osteoblast matrix and mineral deposition. Sclerostin inhibits canonical Wnt/β-catenin signalling, suppressing osteoblast differentiation and bone formation [8, 9], while indirectly promoting bone resorption through osteoblast–osteoclast coupling [10, 11]. Sclerostin expression increases during mechanical unloading, suppressing bone formation and thereby contributing to reduced bone strength [7, 12]. These features make sclerostin a key therapeutic target in postmenopausal osteoporosis. Neutralizing antibodies against sclerostin (Scl-Ab) stimulate bone formation [13] by promoting osteogenic differentiation of progenitor cells and activating bone-lining cells [13-16], while indirectly suppressing osteoclast activity [10, 11]. Clinical studies in postmenopausal women show that monthly romosozumab administration (210 mg) significantly increases bone mineral density (BMD) and reduces vertebral and non-vertebral fracture risk within the first year [6, 14, 15]. Single doses rapidly increase serum P1NP and decrease sCTX, indicating enhanced formation and reduced resorption [17]. However, continued monthly administration beyond six months leads to a partial return of bone formation markers (P1NP, osteocalcin, ALP) toward baseline, suggesting dampened anabolic responses with prolonged exposure [13-15, 18-20]. Treatment-free periods restore bone formation [11, 20, 21], indicating that intermittent dosing, with breaks in antibody administration may preserve efficacy, though the underlying mechanisms are not fully understood.

Mechanistic insights come from preclinical studies in rodents and nonhuman primates. Weekly or twice-weekly Scl-Ab administration in ovariectomized (OVX) rats and cynomolgus monkeys, restores trabecular and cortical bone mass, enhances periosteal and endocortical formation, and improves mechanical strength [10, 19, 22-26]. These studies use higher doses (≥25 mg/kg in rodents) than the clinical equivalent (∼3 mg/kg weekly), due to faster IgG clearance [11, 25, 26]. Prolonged treatment normalizes Wnt target genes and increases secreted antagonists, such as SOST, consistent with feedback regulation observed in humans [11, 20]. Despite these advances, the efficacy of low-dose or intermittent Scl-Ab regimens remains incompletely understood, motivating the exploration of whether lower-exposure schedules can preserve anabolic responses while minimizing feedback dampening.

At the cellular level, osteocytes are key mediators of these effects. Estrogen deficiency reduces osteocyte mechanosensitivity characterized by diminished intracellular calcium oscillations and attenuated responses to mechanical loading[27]. These changes exacerbate osteocyte-mediated osteoclastogenesis and bone resorption [28-30]. Sclerostin inhibition reverses these effects, restoring osteoblast and osteocyte activity via the Wnt/β-catenin signalling pathway. In OVX rats, Scl-Ab treatment upregulates Wnt target and ECM-related genes, while in mice, short-term Scl-Ab exposure increases ECM-associated transcripts such as *Twist1* and *Cgref* [31]. Furthermore, long-term estrogen deficiency alters extracellular matrix gene expression, bone tissue mineral distribution, and the lacunar-canalicular network indicative of perilacunar remodelling and disrupted mechanosensory capacity [32]. Human studies corroborate these findings, showing more heterogeneous bone mineral distribution and increased hypermineralized regions under estrogen deficiency [33]. However, the effectiveness of Scl-Ab treatment during early bone loss, before secondary mineralization occurs, remains unexplored. We hypothesize that early Scl-Ab treatment may prevent secondary mineralization by targeting perilacunar resorption and osteocyte apoptosis in postmenopausal osteoporosis.

The objective of this study was to determine whether initiating monthly low-dose Scl-Ab treatment three weeks post-ovariectomy could: (1) mitigate osteoclastogenesis and bone resorption, (2) preserve perilacunar remodelling, (3) prevent secondary mineralization, and (4) maintain osteocyte mechanosensitivity during estrogen deficiency.

## Methods

### Animal model

Female retired breeder Wistar rats (6 months old, Charles River, Ireland) underwent bilateral ovariectomy (n = 8), confirmed by necropsy. To study subcutaneous delivery of intermittent low-dose Scl-Ab [26] (rodent version of romosozumab: Scl-Ab; r13c7, UCB Pharma, UK / Amgen Inc., UK), ovariectomized rats were assigned to either (a) no treatment (Control, n = 4, body weight: 382 ± 39 g) or (b) Scl-Ab (2 mg/kg/month, n = 4, body weight: 399 ± 49 g), administered every 4 weeks starting 3 weeks post-ovariectomy. This time point corresponds to the initial phase of rapid bone loss but precedes secondary mineralization changes [1]. The 2 mg/kg monthly Scl-Ab regimen was intentionally selected as a low-exposure exploratory dose, lower than the approximate clinical equivalent of 3 mg/kg monthly (210 mg romosozumab in humans). Most preclinical studies use 25–50 mg/kg weekly [11, 25, 26], representing supraphysiologic exposures relative to humans. Our goal was to evaluate osteocyte-specific responses under a conservative, early-intervention dosing schedule, rather than to reproduce maximal pharmacological effects. Scl-Ab was prepared in sterile phosphate-buffered saline (PBS) for subcutaneous injection. Euthanasia occurred 3 weeks after the final injection. Procedures followed guidelines from the University of Galway’s Animal Care and Research Ethics Committee (ACREC) and the Health Products Regulatory Authority (HPRA).

### Calcein Labelling

To label actively mineralizing bone surfaces animals received two intraperitoneal calcein injections (20 mg/kg) at 14 and 4 days before euthanasia. This enabled quantification of mineral apposition rate (MAR) over a 10-day interval. At week 14, vertebrae were isolated from Control (n = 4) and Scl-Ab (n = 4) rats, fixed in 4% paraformaldehyde, and processed for analysis. Trabecular bone within the lumbar vertebral body (L4) was sectioned using a diamond-blade low-speed saw (Isomet™, Buehler, IL, USA), embedded in epoxy, cut into 1.5-mm slices, and mounted on SuperFrost® Plus slides (Menzel Gläser). Calcein-labelled sections were imaged by fluorescence microscopy at ×20 magnification and calibrated (μm/pixel). The total bone surface length (BS) within the trabecular region of interest (ROI) was traced, and the lengths of double-labelled (dL) and single-labelled (sL) surfaces were measured along the same BS using ImageJ. Mineralizing surface per bone surface (MS/BS) was calculated as MS/BS = [(dL + 0.5 × sL) / BS] × 100. Mineral apposition rate (MAR, μm/day) was determined from double labels as the mean inter-label distance divided by the 10-day interval between injections. Bone formation rate per bone surface (BFR/BS) was then calculated as BFR/BS = MAR × (MS/BS fraction). For each animal, measurements were averaged across three sections and at least five fields per section; group means ± SD are reported. All analyses were performed blinded to treatment.

### In vivo and Ex vivo Nano-CT Imaging and Analysis

In vivo micro-CT scanning (VivaCT40, Scanco Medical AG) was conducted to investigate the effect of Scl-Ab on morphometry and mineralization during estrogen deficiency. Scans were performed at day 0, week 4, week 8 and week 14 on the right tibia of Control (n = 4) and Scl-Ab (n = 4) groups at 15 μm resolution. Scans were not performed at week 3 (Scl-Ab treatment onset), baseline bone status at this time point was therefore assumed based on previously published OVX data from our laboratory [1]. The X-ray settings, region of interest, and segmentation threshold were applied according to a previously established protocol [1]. Analyses were conducted on the proximal tibia (trabecular VOI). Microarchitectural parameters (Bone volume fraction (BV/TV), trabecular number (Tb.N), trabecular thickness (Tb.Th), and trabecular spacing (Tb.Sp)) were quantified from the 3D reconstruction of the trabecular volume of interest using evaluation scripts in the Scanco Image Processing Language in the proximal tibia of the treatment groups at each time point.

Ex vivo high-resolution nano-CT scanning (Zeiss Xradia Versa 620) was performed at week 14 on right tibiae from control (n = 4) and Scl-Ab (n = 4) groups. Hydroxyapatite (HA) phantoms (QRM, PTW Dosimetry) were used to calibrate intensity measurements. Scans were conducted using a 0.4X objective at 70 kV, 8.5 W, with an LE3 source filter at an isotropic resolution of 9.14 μm. A 3-mm region below the proximal tibial growth plate was analysed to assess trabecular bone microarchitecture; BV/TV, Tb.N, Tb.Th, and Tb.Sp.

Lacunae and peri-lacunar mineral density were also assessed using ultra-high-resolution scans (1.11 μm voxel size). Trabecular bone sections (0.5 mm thick) were prepared, mounted, and scanned with a 4X objective and a 4-second exposure time with same electrical parameters. Lacunae were segmented using Otsu thresholding quantification of peri-lacunar mineral density and lacunar surface area.

### Histological and Immunohistochemical Analysis

Histological assessments (TRAP, H&E, and MMP14 immunohistochemistry) were performed on distal femoral metaphyseal sections. At week 14, femurs from Control and Scl-Ab rats were fixed in 4% paraformaldehyde. Distal femoral metaphyseal sections (1.5 mm) were cut using a diamond blade saw and decalcified in 10% EDTA for 14 days. Decalcification was confirmed via oxalate and physical probing tests. Samples were rinsed in PBS overnight, processed using a Leica ASP300 tissue processor and embedded in paraffin (Leica EG1150H). Paraffin sections (8 μm) were cut using a Leica RM2235 microtome, mounted on SuperFrost® Plus slides, and stored at room temperature, until staining.

TRAP and H&E staining were performed to visualize osteoclast number and osteocytes within lacunae, respectively. Images were captured using a light microscope (Olympus BX43, Olympus, Japan), and quantitative analysis was conducted on ImageJ.

For immunohistochemistry, paraffin-embedded sections were incubated overnight at 4°C with primary anti-MMP14 antibody (1:100; Abcam, ab38971), followed by a secondary antibody (Goat Anti-Rabbit IgG H&L (HRP); Abcam, ab6721) for 1 hour at room temperature. Slides were developed using DAB substrate chromogen solution for 5 minutes. Control sections, received antibody diluent instead of the primary antibody, were processed alongside experimental samples to confirm specificity. The prevalence of MMP14+ osteocytes, normalized to total bone area, was assessed using ImageJ [34].

### Gene Expression Analysis

Gene expression analysis was carried out on cortical bone isolated from the tibial diaphysis. Tibiae from control (n=4) and Scl-Ab (n=4) groups were dissected, washed with DNase/RNase-free PBS, and cortical bone isolated. Samples were washed to remove bone marrow, snap-frozen in liquid nitrogen, and stored at -80°C. Total RNA was extracted using the RNeasy Fibrous Tissue Mini Kit (Qiagen) and converted to cDNA with the SuperScript VILO Kit (Invitrogen). RNA quality was assessed by 260/280 and 260/230 absorbance ratios. Scl-Ab-treated samples showed acceptable 260/280 ratios (1.8–2.0), with slightly lower 260/230 ratios (<2), but RNA quality was sufficient for mRNA microarray analysis, confirmed by consistent signal intensities and reproducible biological replicates.

TaqMan® microarray analysis was performed using Applied Biosystems TaqMan® Array (format: FAST 96-well) on a StepOnePlus™ PCR system, with Rpl13a as the reference gene. The study examined gene expression related to bone resorption and matrix degradation (RANKL, OPG, Sost, NFATc-1, CtsK, MMP9, MMP13, Ccr2, ATG7, Raptor), apoptosis (Tp53, Casp3, Lamp1), bone formation and matrix production (OPN, Col1α1, Col1α2, OCN, Alp (osteoblast-associated) and DMP1, PHEX, TGFβ1, Klf10 (osteocyte-enriched)), other regulators (Ager, Fn1 (stress- and ECM-associated)) and mechanosensation and mechanotransduction (VCL, TRPV4, ADCY6, AXIN2, ITGβ1, ITGα5, ITGαV, CX43, RYR, PKD1, CAV1). Results are expressed as relative quantitative changes normalized to age-matched OVX controls (Control), with normalization to the housekeeping gene Rpl13a. Eight microarrays were performed, equally divided between Control and Scl-Ab groups.

### Statistical analysis

Statistical analyses were performed using GraphPad Prism. Differences between Control and Scl-Ab groups were assessed via Student’s t-tests, with significance set at p ≤ 0.05. CT parameters (BV/TV, Tb.N, Tb.Th, Tb.Sp) were tested for normality; normally distributed data were analysed with t-tests, while non-normal data were analysed using Mann-Whitney tests. Two-way ANOVA was conducted to assess time and treatment interactions, followed by Tukey’s post hoc tests for significant interactions. Results are presented as mean ± SD. This study was designed as a pilot exploratory experiment with small group sizes (n = 4).

## Results

### Bone formation recovery in Scl-Ab treatment of ovariectomy-induced bone loss

To assess the longitudinal effects of Scl-Ab treatment on bone microarchitecture and mineralization, we performed both in vivo micro-CT and ex vivo nano-CT analyses (Figure 1A). Morphometric analysis by in vivo micro-CT (15 μm) indicated no significant differences in bone volume fraction and trabecular microarchitecture between Scl-Ab and Control groups at baseline (day 0), week 4, week 8 and week 14 (Figures 1B, 1C). However, there was a significant decrease in bone volume fraction and trabecular number and a significant increase in trabecular spacing in week 14 compared to day 0 for both Control and Scl-Ab groups.

**Figure 1.**
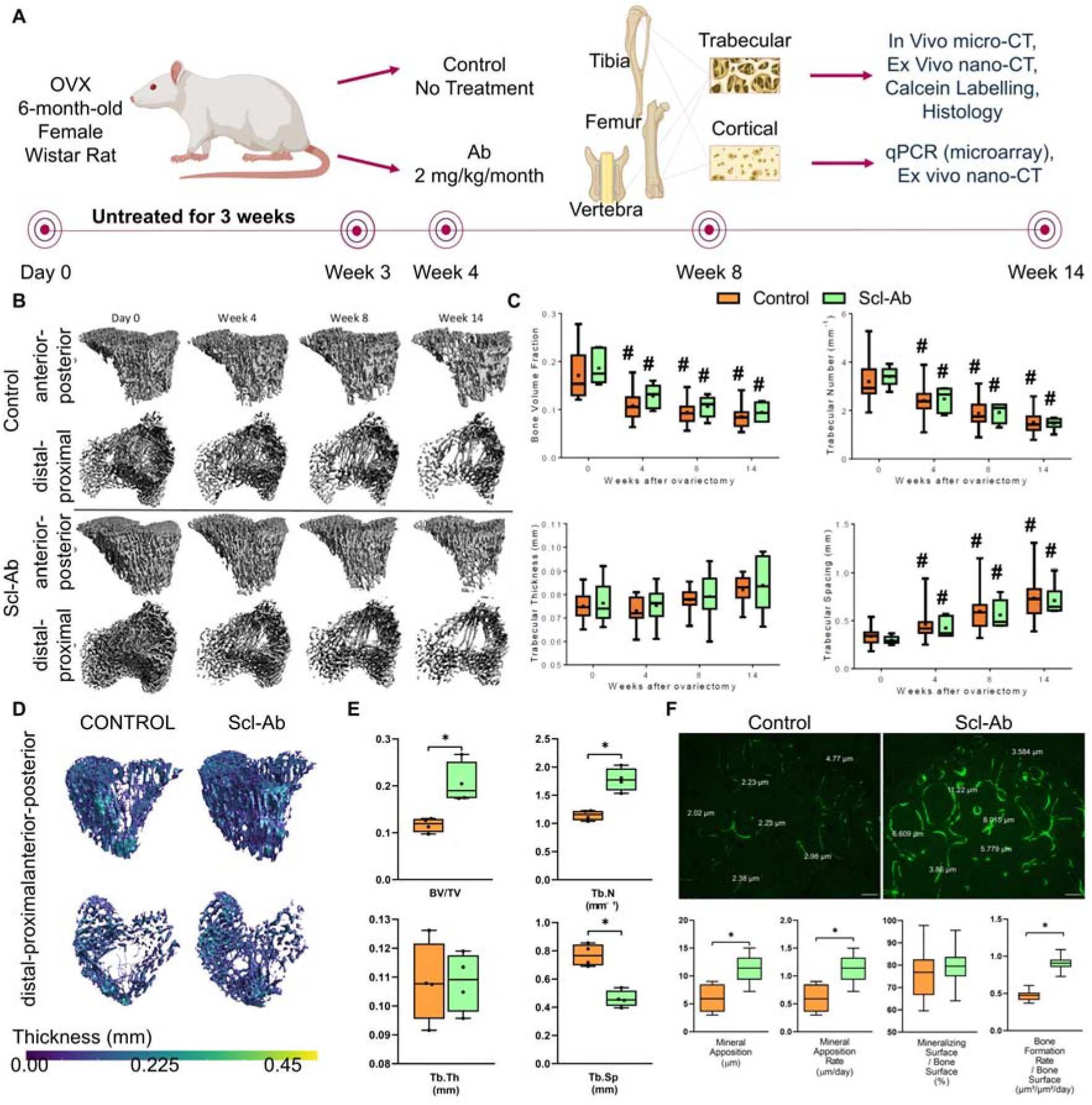
Surface Bone formation in early low-dose Scl-Ab treatment of ovariectomy-induced bone loss. (A) Study design: Female retired breeder Wistar rats were ovariectomized at 6 months. Three weeks post-ovariectomy, the animals were randomly assigned to (i) Control (no treatment) or (ii) Scl-Ab (systemic delivery of 2 mg/kg/month Scl-Ab). (B) 3D micro-CT images of proximal tibial trabecular bone at 0, 4, 8, and 14 weeks post-OVX in both groups, shown in two orthogonal views: anterior-posterior and distal-proximal. (C) Comparison of bone mass and microarchitectural parameters in Control and Scl-Ab groups at 0, 4, 8, and 14 weeks post-OVX. (D) High-resolution 3D nano-CT images of metaphyseal trabecular bone from the proximal tibia of a representative Control and Scl-Ab group at 14 weeks post-OVX, shown in two orthogonal views: anterior-posterior and distal-proximal. (E) Comparison of bone mass and microarchitectural parameters between the Control and Scl-Ab groups 14 weeks post-OVX. (F) Calcein labelling of surface bone formation in Control and Scl-Ab groups, mineral apposition (μm) and rate (μm/day) quantified (n = 4/group). Box plots show median, IQR, and range. *p < 0.05; ^#^p < 0.05 vs. baseline (day 0). Scale = 50 μm. Data in panels B–E from trabecular bone of the proximal tibia; calcein labelling (F) in trabecular bone of a lumbar vertebra (L4).

Morphometric analysis at nano-CT resolution (9.14 μm) revealed significant differences between the Scl-Ab treated and Control groups at week 14 (Figures 1D, 1E). Specifically, there was a significant increase in BV/TV and trabecular number, and a significant decrease in trabecular spacing in the Scl-Ab group compared to the untreated OVX group. No significant differences were observed in trabecular thickness.

Fluorochrome labelling of mineral apposition (Figure 1F) confirmed that Scl-Ab therapy enhanced bone formation in OVX rats. The mineral apposition was 11.54 ± 2.194 μm in the Scl-Ab-treated group compared to 6.16 ± 2.347 μm in untreated OVX controls (p < 0.05). The mineral apposition rate was 1.14 ± 0.22 μm in the Scl-Ab-treated group compared to 0.62 ± 0.23 μm in untreated OVX controls (p < 0.05).

Dynamic histomorphometry revealed that Scl-Ab treatment increased the extent of actively mineralizing surfaces and the overall bone formation rate. Mineralizing surface per bone surface (MS/BS) was higher in Scl-Ab-treated rats (79.6 ± 7.3 %) compared with OVX controls (75.5 ± 10.1 %), although this was not significant. Bone formation rate per bone surface (BFR/BS) was almost doubled in the Scl-Ab group (0.91 ± 0.08 μm^3^/μm^2^/day) relative to controls (0.47 ± 0.06 μm^3^/μm^2^/day; *p* < 0.05). These data confirm that early intermittent Scl-Ab treatment enhances both the rate and extent of trabecular bone formation.

### Osteoclastogenesis and bone resorption are reduced in OVX rats treated early with intermittent low-dose Scl-Ab

Early administration of intermittent low-dose Scl-Ab (2 mg/kg/month) significantly reduced osteoclast activity and gene expression associated with bone resorption. By week 14, TRAP+ staining was significantly lower in Scl-Ab-treated OVX animals than in untreated OVX controls (Figure 2A).

**Figure 2.**
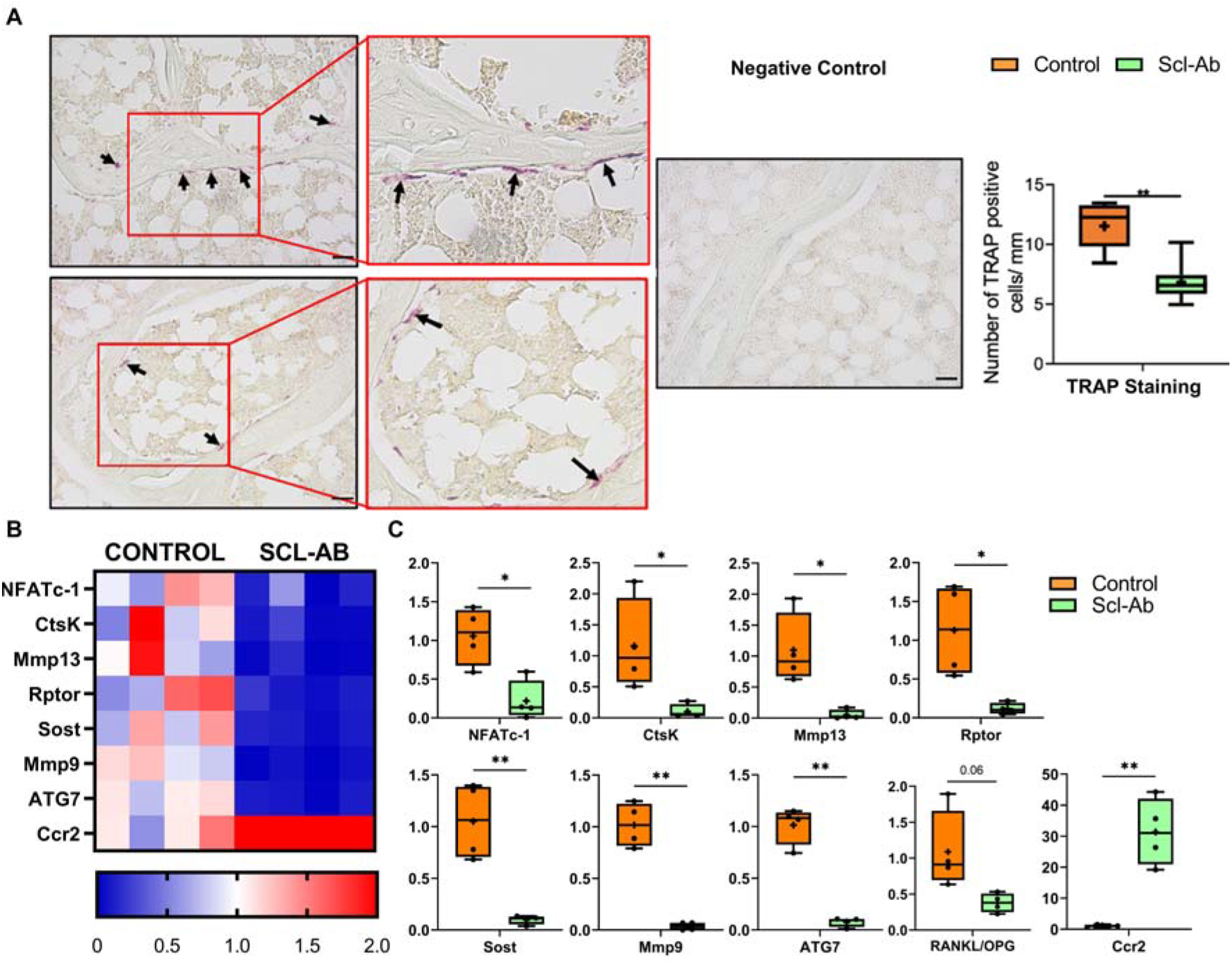
Osteoclastogenesis and bone resorption are reduced in OVX rats treated early with intermittent low-dose Scl-Ab. (A) TRAP+ staining is reduced in OVX animals that received subcutaneous Scl-Ab compared to untreated OVX animals (Control) at week 14. Representative images show TRAP staining (black arrows indicate TRAP+ osteoclasts) and fast green counterstaining of cortical bone in Control and Scl-Ab (magnified images in red boxes). ImageJ analysis was performed to quantify the number of TRAP+ cells per mm. (B) Heat map showing expression of genes involved in catabolic activity and matrix degradation in cortical bone from OVX rats at week 14, comparing untreated controls to those treated with 2 mg/kg/month Scl-Ab starting 3 weeks post-OVX. (C) Scl-Ab treatment significantly lowers expression of Sost, Nfatc1, Ctsk, Mmp9, Ccr2, and Rankl/Opg. *p < 0.05, **p < 0.01; n = 4/group. Scale = 20 μm. TRAP counts and histology are from distal femoral metaphyseal sections (trabecular compartment); gene expression data shown are from cortical bone of the proximal tibial diaphysis.

Gene expression analysis confirmed a downregulation in key bone catabolism markers: NFATc-1 (6-fold), CtsK (12-fold), MMP13, Raptor, Sost (11-fold), MMP9 (25-fold), and ATG7 (10-fold) (Figures 2B, 2C). A trend toward a lower RANKL/OPG ratio was observed in the Scl-Ab group (p = 0.06), suggesting a shift toward reduced osteoclastogenesis. Interestingly, CCR2 expression was upregulated (31-fold) in Scl-Ab-treated OVX animals compared to untreated controls (p < 0.05, Figures 2B, 2C).

### Osteocyte apoptosis and perilacunar remodelling are reduced in OVX rats treated early with intermittent low-dose Scl-Ab

Scl-Ab treatment significantly reduced osteocyte apoptosis in OVX rats. Specifically, the percentage of empty lacunae was significantly lower in Scl-Ab-treated OVX animals compared to untreated OVX controls (p < 0.05, Figure 3A). Apoptosis-related genes were downregulated in Scl-Ab-treated OVX animals: Tp53 (11-fold), Casp3 (11-fold), and Lamp1 (100-fold) (p < 0.01, Figure 3B). Nano-CT revealed no significant difference in mean BMD between groups (*p* = 0.08; Figure 3C). However, treated animals exhibited significantly smaller lacunar surface areas (Figure 3C). The number of MMP14+ osteocytes, indicative of perilacunar remodelling, were significantly reduced in the Scl-Ab-treated group (p < 0.05, Figure 3D). Mineral density significantly reduced near the lacunar edge in Scl-Ab-treated OVX animals compared to controls (*p* < 0.05; Figure 3E). This reduction was localized, diminishing with distance from the lacunae (Figure 3E).

**Figure 3.**
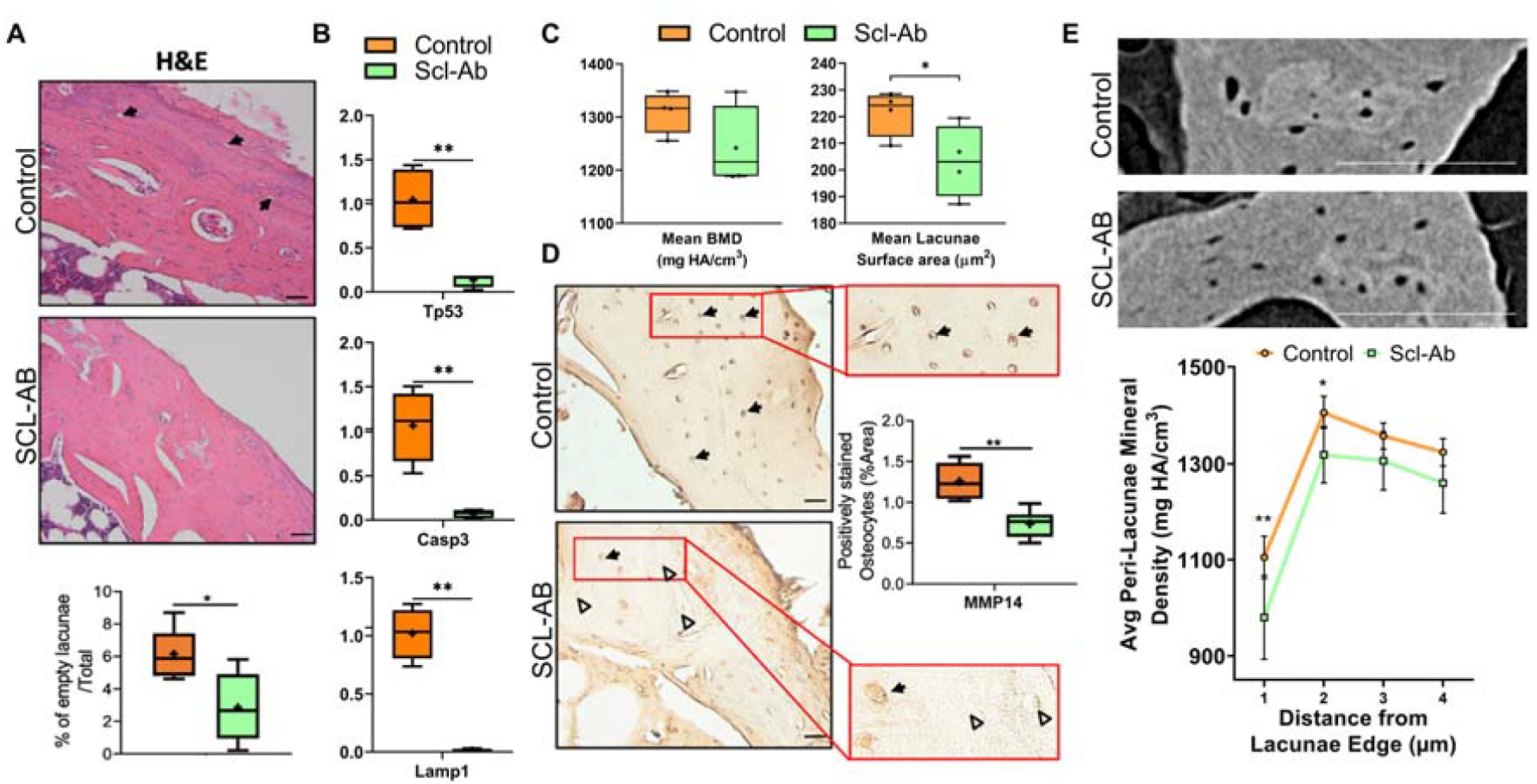
Osteocyte apoptosis and perilacunar remodelling are reduced in OVX rats treated early with intermittent low-dose Scl-Ab. (A) Scl-Ab administration results in a decrease in unoccupied lacunae. Representative images show Haematoxylin and Eosin (H&E) staining of cortical bone from OVX rats at week 14, comparing untreated controls to those treated with 2 mg/kg/month Scl-Ab starting 3 weeks post-OVX. ImageJ analysis was performed to quantify the percentage of empty lacunae (black arrows indicate empty lacunae). (B) Scl-Ab administration results in a decrease in gene expression associated with apoptosis (Tp53, Casp3, and Lamp1). (C) Mean BMD (mg HA/cm^3^), and Mean lacunae surface area (mm^3^) quantified using ultra-high-resolution nano-CT images. (D) Scl-Ab administration results in a decrease in MMP14+ osteocytes. Representative images show MMP14 staining in Control and Scl-Ab (magnified images, red boxes). Black arrows indicate MMP14+ osteocytes, and the white triangle indicates MMP14-osteocytes. ImageJ analysis was performed to assess the prevalence of positively stained osteocytes, normalized to total bone area. (E) Representative nanoCT images of metaphyseal trabecular bone from the proximal tibia from OVX rats at week 14, comparing untreated controls to those treated with 2 mg/kg/month Scl-Ab starting 3 weeks post-OVX, which were evaluated to determine the average peri-lacunae mineral density (mg HA/cm^3^), Scale bar = 20 μm (A and D) and 100 μm (E). *p < 0.05, **p < 0.01. H&E and MMP14 immunohistochemistry are from distal femoral cortical or metaphyseal sections as indicated; nano-CT lacunar/peri-lacunar measures and mean BMD are from cortical VOIs in the proximal tibia.

### Osteocyte gene expression is reduced in OVX rats early treated with intermittent low-dose Scl-Ab

Scl-Ab-treated OVX animals exhibited significant downregulation of several osteocyte-enriched genes, including *Klf10* (6-fold), *DMP1*, and *PHEX* (10-fold each), which are critical for osteocyte-mediated regulation of mineral homeostasis and signalling within the bone matrix. Additionally, *OPN* (osteopontin), a matrix protein expressed by both osteoblasts and osteocytes, was significantly reduced (30-fold), suggesting a broader suppression of mineral-associated signalling within the osteocytic network. Importantly, expression of *AGER*, a receptor known to be upregulated in osteocytes under oxidative or inflammatory stress, was significantly increased (82-fold) in the Scl-Ab-treated group compared to untreated OVX controls (p < 0.05; Figures 4A, 4B), pointing to potential activation of stress response pathways in osteocytes.

**Figure 4.**
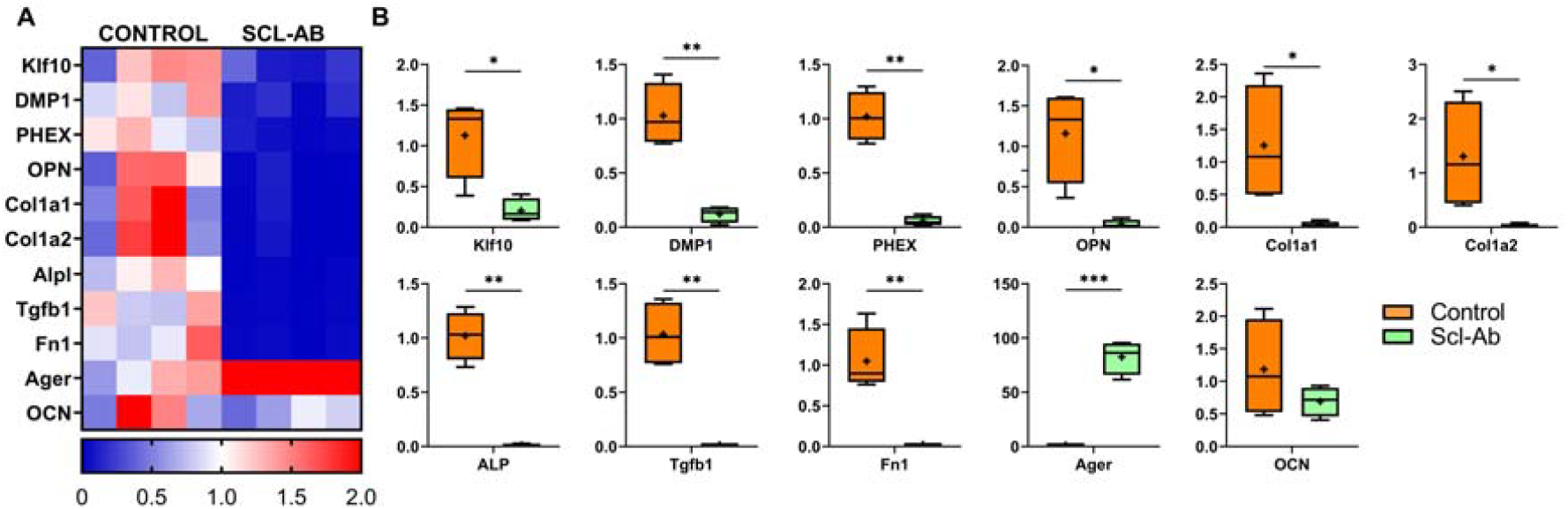
Scl-Ab administration decreases gene expression related to anabolism and bone matrix formation. (A) A heat map displaying anabolic and bone matrix formation gene expression in cortical bone from OVX rats at week 14, comparing untreated controls to those treated with 2 mg/kg/month Scl-Ab starting 3 weeks post-OVX. (B) Boxplots indicate a significant decrease in markers of anabolism and bone matrix formation (OPN, ALP, DMP1, PHEX) following Scl-Ab administration. n = 4 rats/group. *p < 0.05, **p < 0.01, ***p < 0.001. All gene expression data were obtained from cortical bone isolated from the proximal tibia (see Methods).

### Osteoblast gene expression is reduced in OVX rats early treated with intermittent low-dose Scl-Ab

In parallel, we observed suppression of genes primarily associated with osteoblast function. This included downregulation of *ALP* (100-fold), a key enzyme involved in matrix mineralization, as well as *Col1*α*1* (43-fold) and *Col1*α*2* (65-fold), which encode the major structural components of type I collagen in the bone matrix. Additionally, expression of *TGF*β*1* (50-fold) and *Fn1* (55-fold), both involved in osteoblast-mediated matrix signalling and remodelling, was significantly reduced (p < 0.01; Figures 4A, 4B). No significant change was observed in *OCN* (osteocalcin) expression (p = 0.24), a late marker of osteoblast differentiation and bone formation.

### Downregulation in mechanotransduction gene expression in OVX rats early treated with intermittent low-dose Scl-Ab

To explore the impact of Scl-Ab treatment on mechanotransduction pathways, we performed a heatmap analysis of related gene expression, which revealed downregulation in the Scl-Ab-treated OVX group: Vcl (55-fold), Integrin α5 (5-fold), Integrin αV (25-fold), Integrin β1 (50-fold), CX43 (33-fold), Axin2 (11-fold), Adcy6 (10-fold), IFT88 (6-fold), Pkd1 (4-fold), and Cav1 (16-fold) (p < 0.01, Figure 5A, 5B). Conversely, RyR expression, a key regulator of intracellular calcium signalling, was significantly upregulated (22-fold) in Scl-Ab-treated OVX animals (p < 0.05, Figure 5A, 5B). TRPV4 expression remained unchanged (p = 0.41).

**Figure 5.**
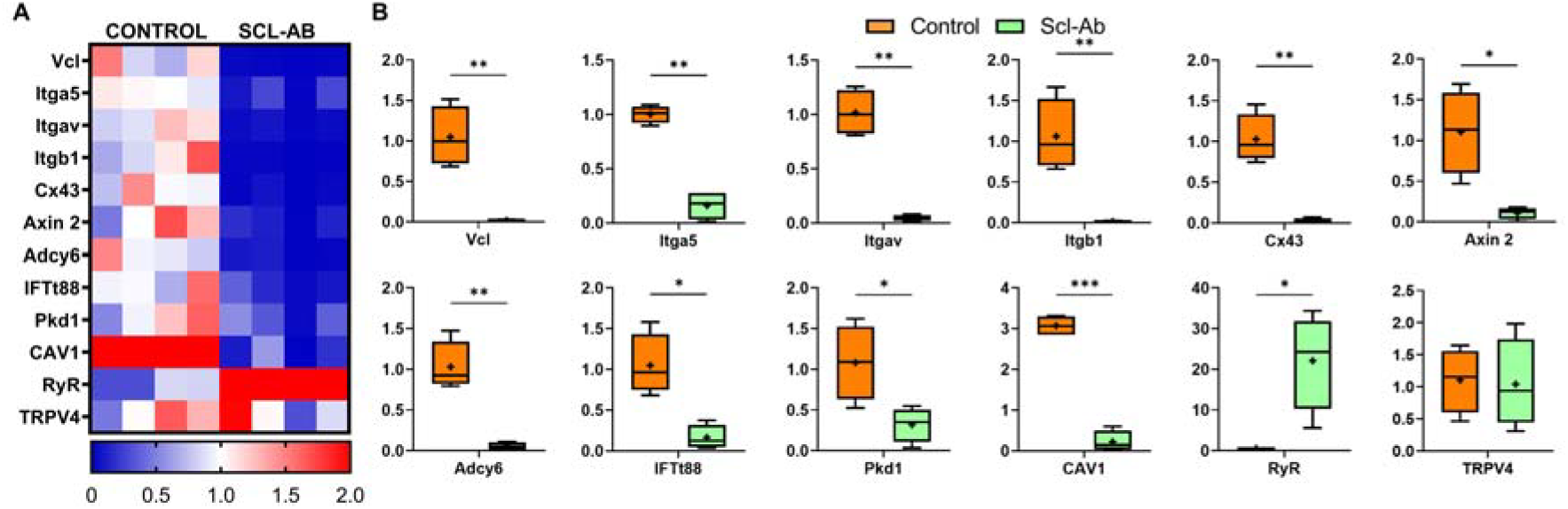
Scl-Ab Administration Downregulates Gene Expression Related to Mechanotransduction and Mechanosensation (A) A heat map displaying gene expression in cortical bone from OVX rats at week 14, comparing untreated controls to those treated with 2 mg/kg/month Scl-Ab starting 3 weeks post-OVX. (B) Boxplots reveal a significant decrease in markers of mechanotransduction and mechanosensation following Scl-Ab administration. n = 4 rats/group. *p < 0.05, **p < 0.01, ***p < 0.001. All gene expression data were obtained from cortical bone isolated from the proximal tibia (see Methods).

## Discussion

The primary objective of this study was to assess the effectiveness of administering intermittent low-dose Sclerostin Antibody (Scl-Ab, 2 mg/kg/month) during the initial phase of rapid bone loss, prior to the onset of secondary mineralization changes [1]. Early administration of intermittent low-dose Scl-Ab reduces osteoclastogenesis and osteocyte apoptosis. It also limits perilacunar remodelling, secondary mineralization, and changes in osteocyte mechanosensitivity associated with estrogen deficiency (see Figure 6). Fluorochrome labelling revealed an increase in bone formation on trabecular surfaces following Scl-Ab therapy in OVX rats. This enhanced bone formation, combined with the suppression of osteoclastogenesis, resulted in an increase in bone volume. These results suggest that early intervention with intermittent low-dose Scl-Ab therapy can mitigate bone loss and osteocyte-mediated secondary mineralization.

**Figure 6.**
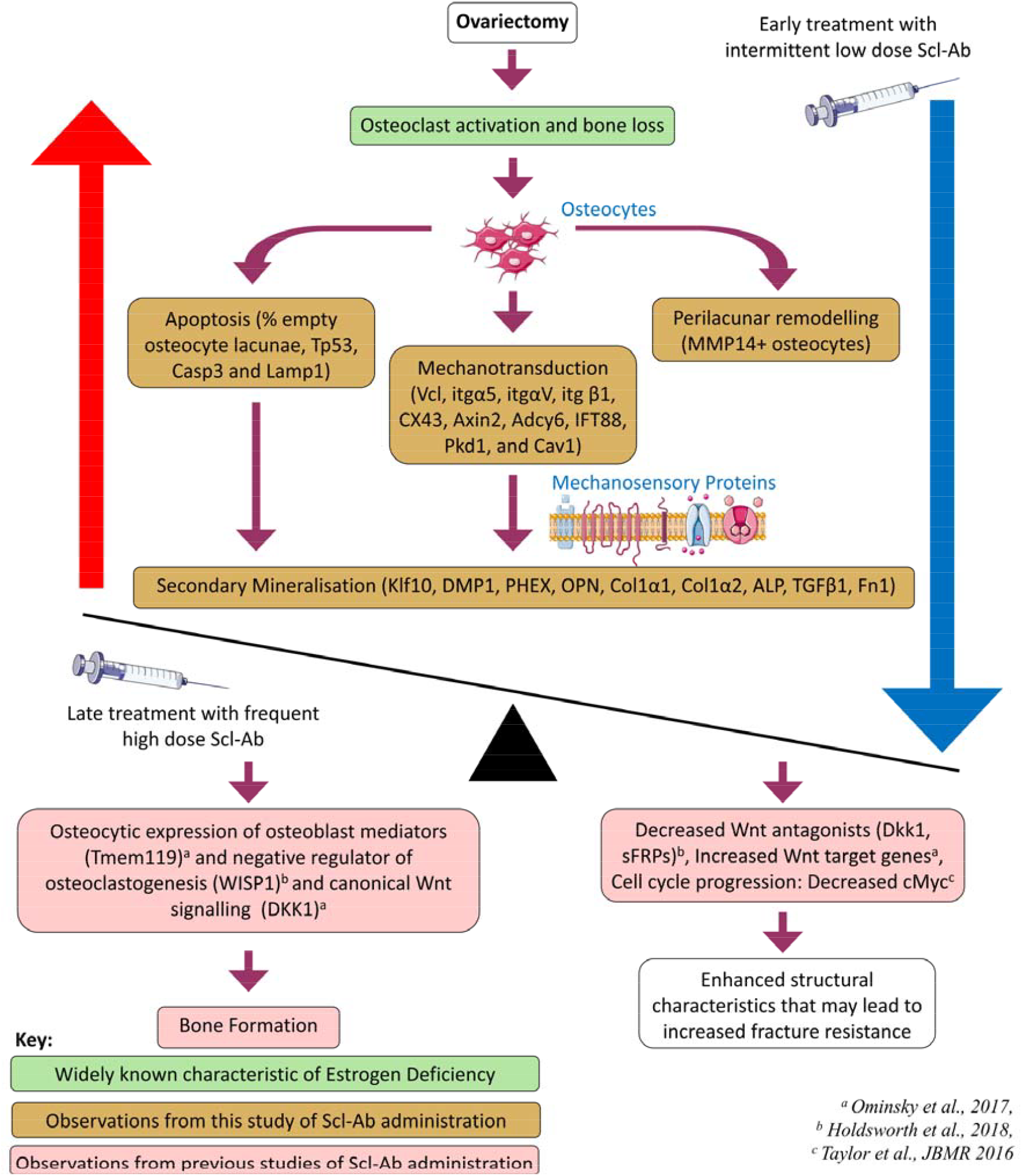
Flowchart depicting the research from this study in the context of a proposed theory of the sequence of events during estrogen deficiency, with early administration of intermittent (monthly) low-dose Scl-Ab antibody and late administration of frequent (weekly) high-dose Scl-Ab.

Several limitations should be considered in this study. Firstly, the estrogen-deficient rat model may not fully replicate human osteoporosis, and individual reproductive histories were unavailable for these retired breeders. Nonetheless, the OVX rat is a widely established model of postmenopausal osteoporosis, because it recapitulates rapid bone loss following estrogen depletion and subsequent remodelling, and reproductive variability exists in the human population. Secondly, the effect of Scl-Ab injections on bone loss was less pronounced at the low dose (2 mg/kg) compared to the higher doses (25 mg/kg) reported to be effective in other OVX rat studies [24]. Additionally, the administration frequency in our study (once per month) differed from other studies, which typically administer Scl-Ab once or twice per week. Despite the lower cumulative exposure, a 2 mg/kg/month dose of Scl-Ab was sufficient to reduce osteoclast number, MMP14 expression in osteocytes, and the incidence of empty lacunae in OVX rats, demonstrating measurable efficacy even under conservative dosing conditions. Fourthly, mechanotransduction effects were inferred from gene expression and were not functionally validated, therefore interpretations regarding osteocyte signalling remain associative. The inclusion of mechanosensitive gene profiling nevertheless provides valuable insight into potential pathways involved in osteocytic responses to Scl-Ab. Finally, the MAR was measured in vertebral trabecular bone, whereas other structural and molecular analyses were performed in the tibia and femur. Given the site-specific nature of bone adaptation to mechanical loading, pharmacological responses to Scl-Ab may differ between vertebral and long bones, and the vertebral MAR may not directly represent mineralization kinetics in the tibia or femur, where strain patterns and remodelling dynamics differ. Also, osteocyte morphology and TRAP[osteoclasts were quantified in trabecular bone, whereas gene expression was analysed in cortical bone. Because remodelling activity and mechanical environments differ between trabecular and cortical compartments, cross-compartment comparisons should be interpreted with caution.

Our findings demonstrate that early administration of intermittent low-dose Scl-Ab in estrogen-deficient rats reduces osteoclastogenesis, consistent with its known dual effects on bone formation and resorption at higher doses [10, 11]. We observed reduced expression of NFATc-1, a key regulator of osteoclast function [35]. Additionally, levels of CtsK (critical for bone resorption) and Raptor (a component of the mTOR pathway) were decreased [36, 37]. Together, these results confirm that early low-dose Scl-Ab reduces both osteoclast formation and activity.

Significant differences in trabecular bone mass and microarchitecture by week 14 align with established effects of Scl-Ab in mitigating bone loss in younger OVX rat models at higher and more frequent doses (25 and 50 mg/kg weekly), especially when administered at later stages (e.g., 8 weeks post-OVX) or in severe osteoporosis with unloading [11, 38]. Notably, in our study, Scl-Ab treatment began at three weeks post-OVX, near the end of the rapid bone loss phase, when 10–30% of bone mass had already been lost [1]. While higher and more frequent regimens (25–50 mg/kg weekly) are known to elicit stronger pharmacodynamic effects in OVX rodents, these doses far exceed clinical exposure levels. The present study used a deliberately conservative monthly dose (2 mg/kg), which is lower than the approximate human-equivalent exposure (∼3 mg/kg/month, 210 mg romosozumab). This strategy aimed to assess osteocyte and bone-remodelling sensitivity under low-exposure conditions that better mimic early-intervention therapy rather than pharmacological saturation. The modest response observed likely reflects this conservative dosing but offers valuable insight into threshold-level efficacy and adaptive responses to intermittent sclerostin inhibition.

We report that early intermittent low-dose Scl-Ab treatment also reduces osteocyte apoptosis and the prevalence of empty lacunae, consistent with prior findings in glucocorticoid-induced osteoporosis models [39]. Moreover, Scl-Ab treatment led to reduced expression of apoptosis markers (Tp53, Casp3) and decreased lysosomal activity (Lamp1), suggesting a concurrent suppression of osteocyte death and its protective cellular responses. We also observed inhibition of MMP14-dependent perilacunar remodelling.

Micropetrosis, the mineral infilling of empty lacunae, commonly follows osteocyte apoptosis, whereas perilacunar remodelling can degrade mineral surrounding osteocytes. Osteocyte loss is typically associated with smaller lacunae, while perilacunar remodelling may cause lacunar expansion. In our study, Scl-Ab treatment led to a decrease in lacunar surface area and a reduction in mineral density within the 2-micron region surrounding lacunae in OVX animals. Further research is required to elucidate how Scl-Ab balances suppression of micropetrosis and perilacunar remodelling, and how this affects osteocyte regulation of their local microenvironment [7, 12].

Ovariectomy has previously been shown to increase osteocyte canalicular diameter [3], potentially enhancing fluid shear stress and making osteocytes more susceptible to apoptosis [3, 40-42]. In this study, Scl-Ab treatment reduced apoptosis markers (Tp53, Casp3) and lysosomal activity (Lamp1), supporting a protective mechanism against osteocyte death. Interestingly, a prior study in rats reported increased p53 expression in osteocytes following Scl-Ab treatment [11], highlighting the potential for context-dependent or time-sensitive effects of Scl-Ab on osteocyte stress responses. Adding further complexity, we observed upregulation of AGER, a receptor implicated in cellular stress, after early intermittent low-dose Scl-Ab treatment, suggesting a nuanced and multifaceted role for Scl-Ab in osteocyte biology. Elevated AGER expression may reflect a compensatory response [43] to microenvironmental changes in local mineral or matrix conditions induced by early intermittent low-dose Scl-Ab treatment, potentially indicating transient signalling linked to bone remodelling or a persistent stress response with functional implications.

We also observed a significant increase in mineral apposition rate, indicating enhanced osteoblast-driven surface bone formation in Scl-Ab-treated animals. Although early osteogenic genes (ALP, Col1α1, Col1α2) were downregulated at week 14, the bone formation observed likely occurred earlier in treatment, driven by initial osteoblast activity. We propose that the discrepancy between increased trabecular MAR in the vertebra and reduced osteoblast gene expression in cortical bone may reflect both temporal and compartmental differences in Scl-Ab action: whereby an early, robust anabolic response occurs in trabecular regions, which followed by later adaptation in cortical bone. Trabecular bone has a higher surface-to-volume ratio, greater osteoblast and progenitor density, and faster remodelling kinetics compared with cortical bone, facilitating rapid osteoblast recruitment and bone formation. This pattern is consistent with transient osteoprogenitor proliferation described in prior Scl-Ab studies [44, 45]. In aged OVX rats, Scl-Ab initially promotes progenitor proliferation, leading to rapid osteoblast recruitment. Although proliferation declines over time, bone formation remains elevated due to enhanced osteoblast efficiency [46, 47]. These dynamics differ from those seen with high-dose or delayed Scl-Ab regimens, which tend to more robustly upregulate osteogenic gene expression at later disease stages [11, 24].

Importantly, while surface bone formation increased, osteocyte-mediated secondary mineralization was reduced. To our knowledge, this is the first report showing downregulation of osteocyte-enriched genes (Klf10, DMP1, and PHEX) following early intermittent low-dose Scl-Ab treatment. We also found downregulation of mechanotransduction-related genes (ItgαV, Itgβ1, and Cav1), which are essential for osteocyte mechanosensation and Wnt signalling. Short-term Scl-Ab treatment has also been associated with reduced osteocyte mechanosensitivity, possibly due to the presence of immature osteocytes following new bone formation. However, longer-term studies show a return to normal osteocyte signalling, which may explain the eventual reduction in bone formation [48]. We propose that Scl-Ab may initially reduce osteocyte sensitivity to mechanical loading, a potential protective adaptation to the altered mechanical environment following bone loss. By dampening mechanotransduction, Scl-Ab may help mitigate stress-induced osteocyte apoptosis and pathological mineralization [49]. However, these findings are correlative, we assessed gene expression changes in cortical bone but did not perform functional loading experiments. Therefore, mechanistic interpretations will require validation in future studies using controlled mechanical loading, combined with measurements of early mechanosignalling events (e.g., calcium transients, immediate early gene activation) and subsequent bone formation outcomes.

In summary, early intermittent low-dose Scl-Ab treatment preserves osteocyte viability and prevents perilacunar remodelling, while suppressing bone resorption and promoting surface bone formation. Moreover, low-dose treatment can prevent osteocyte-driven secondary mineralization and mechanosensitivity through distinct cellular mechanisms.

## Statement on conflicts or disclosures

Syeda Masooma Naqvi, Wahaaj Ali, Hollie Allison, Laura M. O’Sullivan, Juan Alberto Panadero-Perez, Jessica Schiavi-Tritz and Laoise M. McNamara declare the following which may be considered as potential competing interests: The authors report sclerostin antibody was provided by UCB Ltd. Gill Holdsworth reports a relationship with UCB Ltd that includes employment.

## Acknowledgements

This publication has emerged from research supported by SFI, co-funded under the European Regional Development Fund (14/IA/2884), and the ERC Consolidator Grant (MEMETic: 863795). The authors would like to acknowledge Servier Medical Art for their image bank, which was used to produce the figures, and UCB Pharma and Amgen Inc. for providing the Scl-Ab via a materials transfer agreement.

---

**Supplementary Figure 1.**
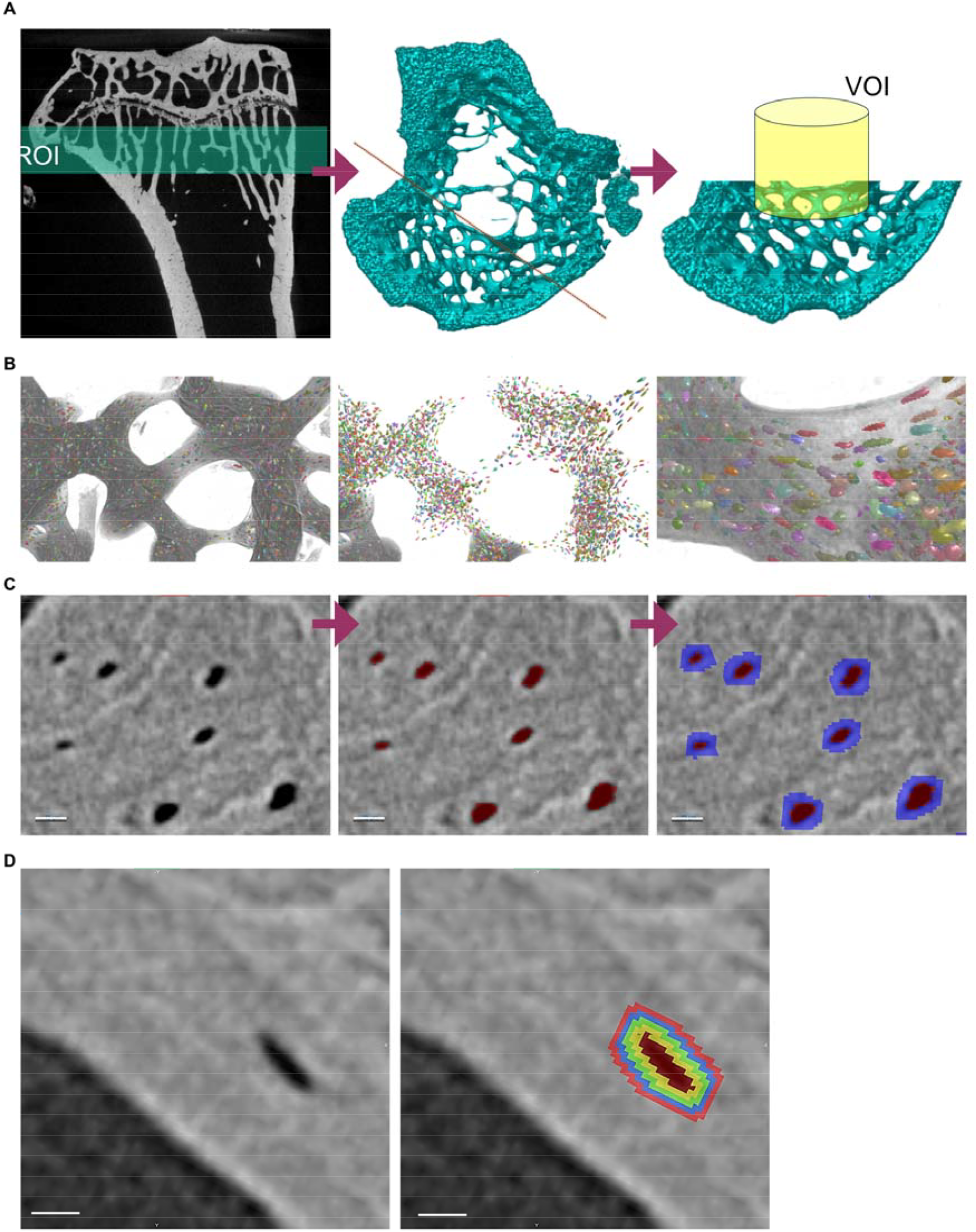
(A) Sample preparation for nano-CT imaging. A ∼0.5 mm slice was extracted from the whole bone using a diamond blade. (B) A rendered VOI from a representative sample, with segmentation and labelling of individual trabecular structures. (C) Peri-lacunar mineral density assessment. Lacunae ROIs were dilated by 2 pixels, with the inner lacunar volume removed to isolate the peri-lacunar region for each lacuna. Lacunae are segmented in red, and peri-lacunar regions are segmented in blue. (D) Peri-lacunar mineral density variation. A similar strategy was used to analyse peri-lacunar mineral variation. New ROIs were defined by dilating the original ROI by 1 pixel at 1.1-micron spacing. Lacunae are segmented in red, with peri-lacunar regions visualized in sequentially expanded layers.

